# Latent biophysical diffusion properties underlying cerebral microstructure revealed through multimodal MRI covariation in the squirrel monkey

**DOI:** 10.64898/2026.01.19.700428

**Authors:** Cláudio Eduardo Corrêa Teixeira, Liliane Almeida Carneiro, Aline Amaral Imbeloni, Pedro Fernando da Costa Vasconcelos

## Abstract

Diffusion MRI provides multiple quantitative descriptors of neural tissue derived from distinct physical and biophysical models, each probing different aspects of cerebral microstructure. While these metrics are often interpreted independently, their joint statistical structure may encode information about latent biophysical properties that constrain and organize brain microarchitecture. Here, we investigate whether stable patterns of multimodal diffusion MRI covariation reveal underlying biophysical diffusion properties of cerebral microstructure in the squirrel monkey (*Saimiri sciureus*). Using a high-resolution multimodal diffusion MRI dataset acquired at 11.7T (400 μm isotropic resolution) from 15 adult subjects, we analyzed tensor-based metrics alongside advanced multicompartment and axonal models, including NODDI and ActiveAx. Voxelwise and region-level correlation analyses were performed across multiple spatial scales to examine the structure, redundancy, and stability of relationships among diffusion-derived maps. We show that metrics originating from conceptually distinct models consistently organize into a low-dimensional structure characterized by stable clusters of covariation, robust to changes in parcellation and spatial aggregation. These patterns cannot be explained by trivial metric redundancy alone, but instead suggest convergence toward a reduced set of effective biophysical degrees of freedom governing diffusion behavior in neural tissue, including axonal organization, neurite density, orientation dispersion, and isotropic diffusion components. We interpret these covariation structures as manifestations of latent biophysical diffusion properties—emergent tissue states that are not directly observable through any single metric but become apparent through their structured relationships. From a systems neuroscience perspective, the stability of these latent dimensions supports the view that adult cerebral microstructure reflects the steady-state outcome of neurodevelopmental dynamics operating under biophysical constraints. Rather than providing a regional atlas, this work proposes a conceptual framework for interpreting multimodal diffusion MRI as a projection of an underlying low-dimensional biophysical state space organizing cerebral microstructure in primate brains.

**Key points:**

- Multimodal diffusion MRI metrics derived from distinct biophysical models exhibit stable and non-trivial patterns of covariation, revealing a low-dimensional organization of cerebral microstructure.
- These covariation structures suggest the existence of latent biophysical diffusion properties that are not directly observable through individual metrics but emerge from their structured relationships across spatial scales.
- The stability of these latent dimensions supports the interpretation of adult cerebral microstructure as a structurally stable outcome of neurodevelopmental dynamics governed by biophysical constraints.

## Introduction

The cerebral microstructure of primates emerges from a complex interplay between neurodevelopmental programs, geometric constraints, and biophysical properties of neural tissue. At the macroscopic level, this organization is reflected in conserved patterns of cortical folding, laminar differentiation, and long-range connectivity. At the microscopic and mesoscopic scales, however, cerebral architecture is governed by the arrangement, density, orientation, and packing of neurites, glial processes, and extracellular compartments, whose combined properties constrain diffusion processes within neural tissue (Innocenti & Price, 2005; Budday et al., 2015).

Diffusion magnetic resonance imaging (dMRI) provides a non-invasive window into these microstructural features by probing the interaction between water diffusion and tissue architecture (Le Bihan et al., 2001). Over the past decades, a growing diversity of diffusion models has been developed, ranging from the diffusion tensor model (Basser et al., 1994) to more advanced multicompartment and axonal frameworks, such as NODDI (Zhang et al., 2012) and ActiveAx (Alexander et al., 2010). Each of these approaches captures distinct aspects of microstructure, including anisotropy, neurite density, orientation dispersion, isotropic compartments, and apparent axonal diameter. Nevertheless, diffusion-derived metrics are typically interpreted independently, often leading to fragmented or model-specific descriptions of cerebral microarchitecture (Jones et al., 2013).

An unresolved question in systems neuroscience is whether the joint organization of diffusion metrics contains information beyond that conveyed by individual parameters. Specifically, it remains unclear whether consistent patterns of covariation across diffusion modalities merely reflect mathematical redundancy between models, or instead reveal latent biophysical properties that constrain and organize cerebral microstructure across spatial scales (Novikov et al., 2018). Addressing this question is essential for moving from descriptive mapping toward a more integrative, biophysically grounded interpretation of diffusion MRI data.

Non-human primates offer a particularly advantageous framework for investigating this issue. Their cerebral organization shares key architectural and developmental features with the human brain, while allowing high-resolution acquisitions that are often impractical in vivo in humans (Phillips et al., 2014). In this context, high-field multimodal diffusion MRI datasets acquired in squirrel monkeys (*Saimiri sciureus*) provide an opportunity to examine microstructural organization at submillimetric resolution, enabling detailed analysis of diffusion-derived metrics across cortical and subcortical structures.

Recent multimodal analysis of the *Saimiri* brain suggest not only coherent microstructural gradients across cortex and white matter, but also robust and reproducible patterns of covariation among diffusion metrics derived from distinct biophysical models. Correlation analyses performed at both voxelwise and region-of-interest levels demonstrate that these metrics consistently organize into a small number of stable clusters, largely independent of anatomical parcellation scale. Metrics associated with axonal coherence and intra-neuritic organization tend to covary strongly, while parameters sensitive to isotropic or dispersive diffusion form a complementary cluster. The persistence of this structure across spatial resolutions suggests that these relationships are not driven by local anatomical idiosyncrasies or arbitrary model overlap, but instead reflect underlying constraints imposed by tissue organization.

These observations raise the possibility that multimodal diffusion MRI reflects a reduced set of latent biophysical diffusion properties underlying cerebral microstructure. In this view, individual diffusion metrics represent different projections of a lower-dimensional biophysical state space governed by constraints such as tissue geometry, compartmentalization, and material properties (Callaghan, 1991; Novikov et al., 2019). From a developmental perspective, such latent dimensions may correspond to stable outcomes of neurodevelopmental dynamics, whereby evolving cellular and axonal architectures converge toward structurally stable configurations in the adult brain (Van Essen, 1997; Budday et al., 2015).

In the present study, we build on these findings to examine whether multimodal diffusion MRI covariation can be interpreted as evidence of latent biophysical diffusion properties underlying cerebral microstructure in the squirrel monkey. Rather than providing an exhaustive regional atlas, we adopt a systems-level perspective focused on the structure, stability, and interpretability of intermodal relationships. By framing diffusion MRI metrics as coupled observables of an underlying biophysical organization, this work aims to advance a conceptual framework for integrating multimodal diffusion data into a coherent description of cerebral microarchitecture in primate brains.

## Materials and Methods

### Dataset and Ethical Considerations

This study is based on a publicly available high-resolution multimodal diffusion MRI dataset acquired from 15 adult female squirrel monkeys (*Saimiri sciureus*). The data were obtained using a Bruker BioSpec 11.7T MRI system with isotropic spatial resolution of 400 μm and a multishell diffusion scheme (b = 300, 1000, and 2000 s/mm^2^). The dataset was selected due to its exceptional spatial resolution and signal quality, enabling detailed characterization of cerebral microstructure at mesoscopic scales. The data were made available by the Disconnectome Project under a CC0 1.0 public domain license and are accessible at http://saimiri.bcblab.com/ (Orset et al., 2023). All acquisition procedures were approved by the relevant institutional ethics committee (CEEA Charles Darwin; APAFIS #21086-2019061415485300), and no new animal experiments were conducted for the present study.

### MRI Acquisition Protocol

Diffusion-weighted images were acquired using a 3D segmented EPI sequence (TR/TE = 200/24 ms), with a matrix size of 160 × 160 × 128 and a field of view of 64 × 64 × 51.2 mm^3^, yielding an isotropic voxel size of 0.4 mm. A total of 100 non-collinear diffusion directions were distributed across the multishell scheme. The total acquisition time was approximately 3 hours per subject. B0 images and corresponding b-values and b-vectors were provided as part of the dataset in standardized format.

### Preprocessing and Diffusion Modeling

Initial preprocessing included denoising, reorientation, and correction for motion and eddy-current–induced distortions using established tools (ExploreDTI and FSL eddy). Brain masks were generated and manually refined using BrainSuite. All diffusion volumes were then spatially normalized to a common anatomical space using nonlinear registration (SyN) implemented in ANTsPy. Diffusion models were fitted to the preprocessed data to derive a comprehensive set of microstructural metrics: Diffusion Tensor Imaging (DTI): fractional anisotropy (FA) and mean diffusivity (MD); NODDI: intra-cellular volume fraction (ICVF), orientation dispersion (OD), and isotropic volume fraction (IsoVF); ActiveAx: axonal density (AAx_Dens), volume intra-cellular fraction estimated by ActiveAx (AAx_vICVF), and apparent axonal diameter (AAx_ADiam). These metrics were chosen to represent complementary aspects of neural microstructure, including axonal coherence, neurite density, orientational complexity, isotropic diffusion components, and axonal caliber.

### Computational Environment and Reproducibility

All analyses were conducted offline in a fully reproducible Python-based environment using Google Colab as a standardized Linux platform. The processing pipeline was implemented in Python 3.10 and relied exclusively on open-source libraries, including DIPY, ANTsPy, Nibabel, NumPy, Pandas, SciPy, and Matplotlib. All steps—from data loading to figure generation—were organized into sequential code cells, enabling full replication of the analyses without reliance on external APIs or proprietary software.

### Atlas Construction and Variability Mapping

For each diffusion metric, subject-level maps were aligned to the common template space and averaged to generate population mean maps. Voxelwise standard deviation (STD) maps were computed across subjects to quantify interindividual microstructural variability. To identify regions of heightened variability, voxels exceeding the 95th percentile of the STD distribution for each metric were classified as hotspots of microstructural dispersion. These hotspot maps were used for comparative visualization across modalities but were not the primary focus of the present analysis.

### Multimodal Covariation Analysis

To investigate relationships among diffusion-derived metrics, two complementary correlation analyses were performed: voxelwise correlation analysis and region-based correlation analysis. Correlation analyses were exploratory and descriptive in nature, without formal hypothesis testing.

In the voxelwise correlation analysis, values from all eight metrics were extracted voxel by voxel within the brain mask of the population template. Pearson correlation coefficients were computed for all metric pairs, yielding a symmetric correlation matrix representing global intermodal relationships.

In the region-based correlation analysis, k-means clustering was applied to the anatomical space to generate parcellations with varying granularity (k = 5, 8, 12, and 20 regions) to assess the robustness of intermodal relationships across spatial scales. Mean metric values were extracted within each region, and Pearson correlation matrices were computed for each parcellation. Hierarchical clustering and dendrograms were generated using Euclidean distance and average linkage to examine the stability of intermodal groupings across scales

## Results

In the present study, we interpret their structured covariation as reflecting underlying biophysical diffusion properties of neural tissue. These properties are conceptualized as latent dimensions that constrain diffusion behavior and emerge consistently across spatial scales and modeling approaches. No causal inferences regarding cellular mechanisms are made; instead, the analysis focuses on the stability, organization, and interpretability of intermodal relationships as evidence of a reduced biophysical state space underlying cerebral microstructure.

### Multimodal diffusion metrics reveal complementary but overlapping representations of cerebral microstructure

Figure 1 presents axial, coronal, and sagittal central slices from the population-averaged maps of eight diffusion-derived metrics: fractional anisotropy (FA), mean diffusivity (MD), intra-cellular volume fraction (ICVF), orientation dispersion (OD), isotropic volume fraction (IsoVF), axonal density (AAx_Dens), intra-cellular volume fraction estimated by ActiveAx (AAx_vICVF), and apparent axonal diameter (AAx_ADiam). Despite being derived from the same diffusion-weighted signal, these metrics display markedly distinct spatial contrasts, emphasizing different aspects of cerebral microstructure.

**Figure 1.**
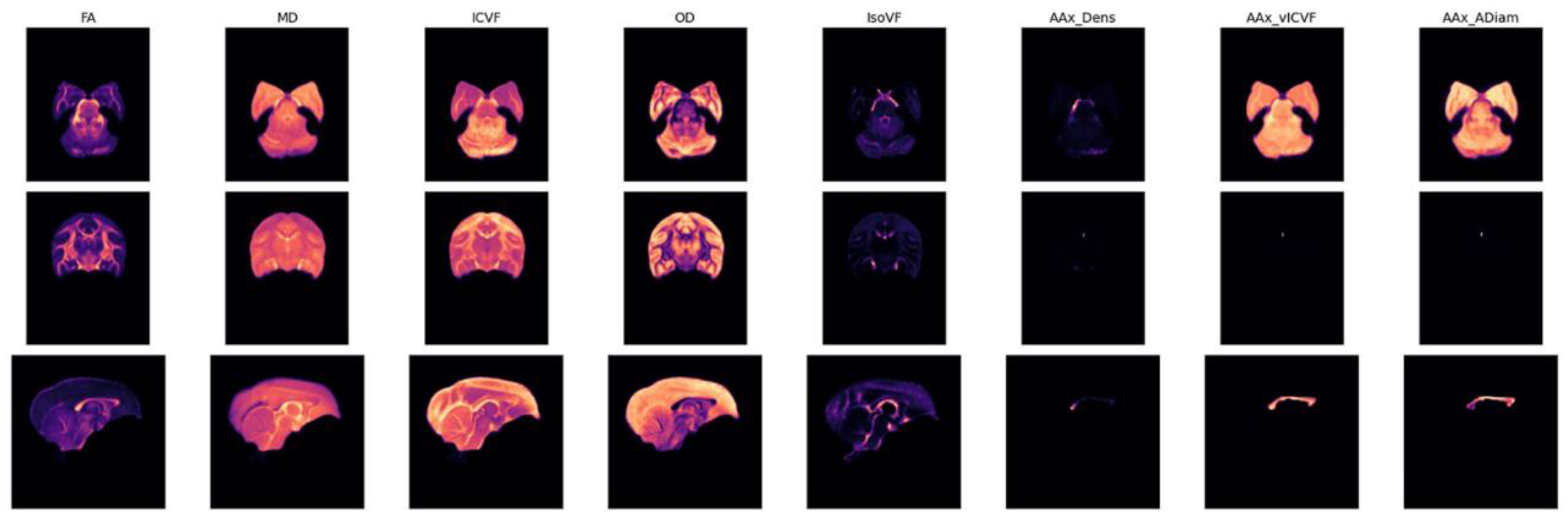
Multimodal diffusion MRI representations of cerebral microstructure in the squirrel monkey. Population-averaged maps of eight diffusion-derived metrics are shown for central axial, coronal, and sagittal slices. Metrics include fractional anisotropy (FA), mean diffusivity (MD), intra-cellular volume fraction (ICVF), orientation dispersion (OD), isotropic volume fraction (IsoVF), axonal density (AAx_Dens), intra-cellular volume fraction estimated by ActiveAx (AAx_vICVF), and apparent axonal diameter (AAx_ADiam). Distinct spatial contrasts across modalities reflect differential sensitivity to orientational coherence, diffusivity, neurite density, dispersion, isotropic components, and axonal-specific properties, highlighting the complementary nature of multimodal diffusion MRI.

Tensor-based metrics highlight global organizational features: FA emphasizes highly coherent fiber bundles and compact white matter structures, while MD shows elevated values in regions with less restricted diffusion, including gray matter and cerebrospinal fluid-adjacent compartments. NODDI-derived metrics further differentiate microstructural components, with ICVF highlighting regions of high neurite density, OD emphasizing areas with complex fiber orientation or dispersion, and IsoVF delineating regions dominated by isotropic diffusion components.

ActiveAx-derived parameters show a more restricted spatial distribution, with signal predominantly localized to major white matter tracts. AAx_Dens and AAx_vICVF exhibit similar anatomical patterns, whereas AAx_ADiam highlights a subset of these regions, reflecting sensitivity to axonal caliber rather than density alone. Notably, although these metrics differ in contrast and spatial extent, their maps consistently delineate common anatomical structures, indicating that they interrogate shared microstructural substrates through distinct biophysical sensitivities.

Collectively, Figure 1 demonstrates that no single diffusion metric provides a complete description of cerebral microstructure. Instead, the multimodal set forms a heterogeneous yet anatomically coherent representation space, motivating the examination of their joint statistical organization.

### Voxelwise covariation reveals structured relationships among diffusion metrics

Voxelwise Pearson correlation analysis across the entire brain reveals a highly structured pattern of intermodal relationships (Figure 2). Several strong positive correlations are observed among metrics derived from distinct modeling frameworks. FA shows moderate correlations with MD (r = 0.43) and ICVF (r = 0.58), indicating partial overlap between orientational coherence, overall diffusivity, and intra-neuritic content. MD correlates strongly with ICVF (r = 0.63) and OD (r = 0.61), suggesting sensitivity to combined effects of neurite density and orientational dispersion.

**Figure 2.**
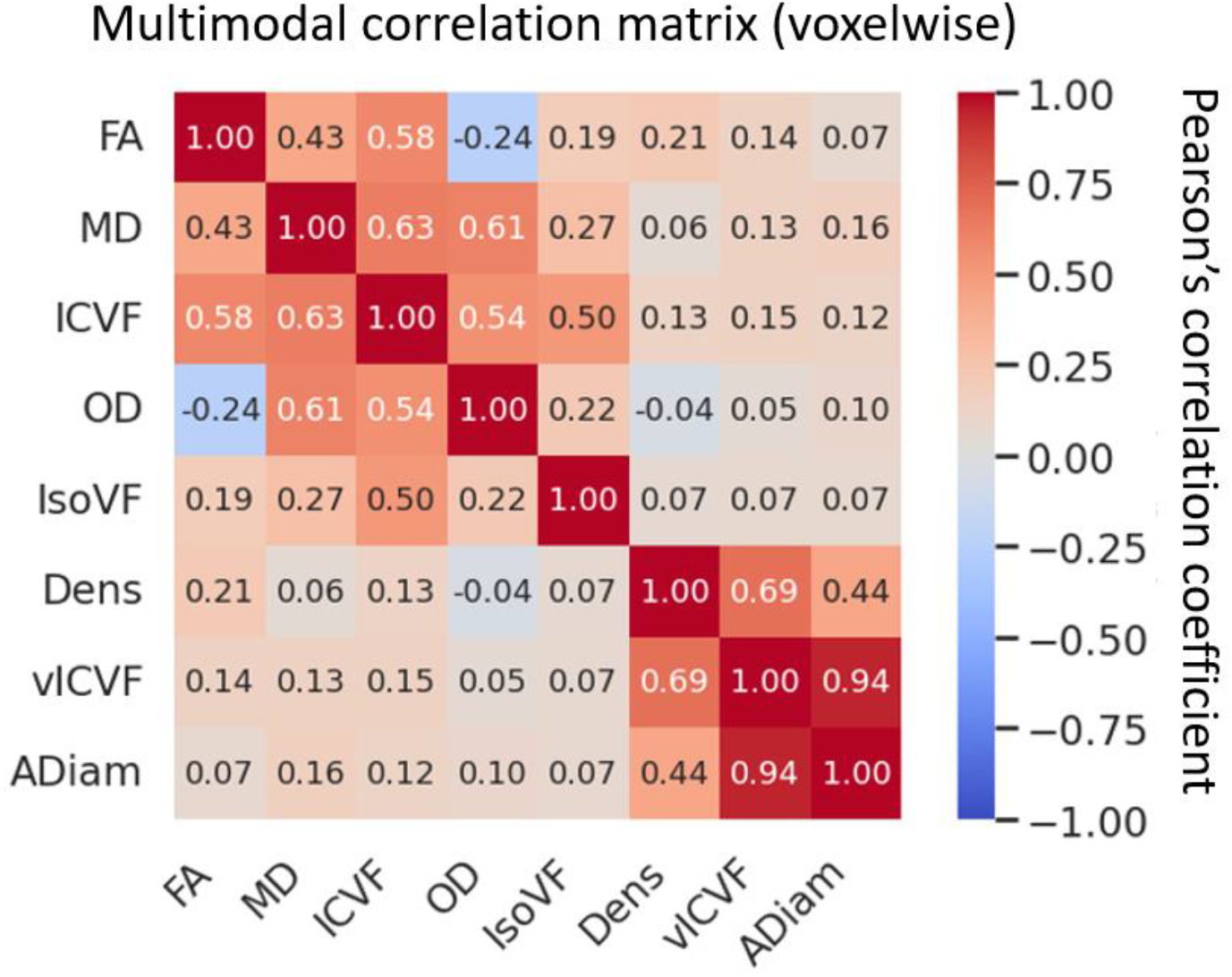
Voxelwise multimodal correlation structure among diffusion MRI metrics. Pearson correlation matrix computed voxelwise across the whole brain shows structured relationships among diffusion-derived metrics, including FA, MD, ICVF, OD, IsoVF, AAx_Dens, AAx_vICVF, and AAx_ADiam. Metrics derived from distinct diffusion models exhibit non-random covariation, with strong positive correlations among axonal and intra-neuritic measures and weaker or antagonistic relationships involving dispersion and isotropic components. The color scale represents Pearson’s correlation coefficients, ranging from −1 to 1.

A pronounced correlation is observed between ActiveAx-derived AAx_vICVF and AAx_ADiam (r = 0.94), reflecting shared sensitivity to axonal-specific features within white matter. AAx_Dens also correlates strongly with AAx_vICVF (r = 0.69), forming a coherent axonal cluster distinct from tensor- and NODDI-based measures. In contrast, IsoVF exhibits generally weak correlations with most metrics, consistent with its sensitivity to a distinct isotropic diffusion component.

Importantly, several negative or near-zero correlations are observed, such as between FA and OD (r = −0.24), indicating that orientational dispersion and anisotropy capture opposing aspects of microstructural organization. Together, these findings demonstrate that diffusion metrics do not vary independently, but instead exhibit structured, non-random covariation patterns across the brain.

### Multimodal covariation patterns persist across spatial scales

To assess the robustness of voxelwise relationships, diffusion metrics were averaged within regions defined by k-means clustering at multiple spatial scales (k = 5, 8, 12, and 20). As shown in Figure 3, the overall structure of intermodal correlations is preserved across parcellation granularities.

**Figure 3.**
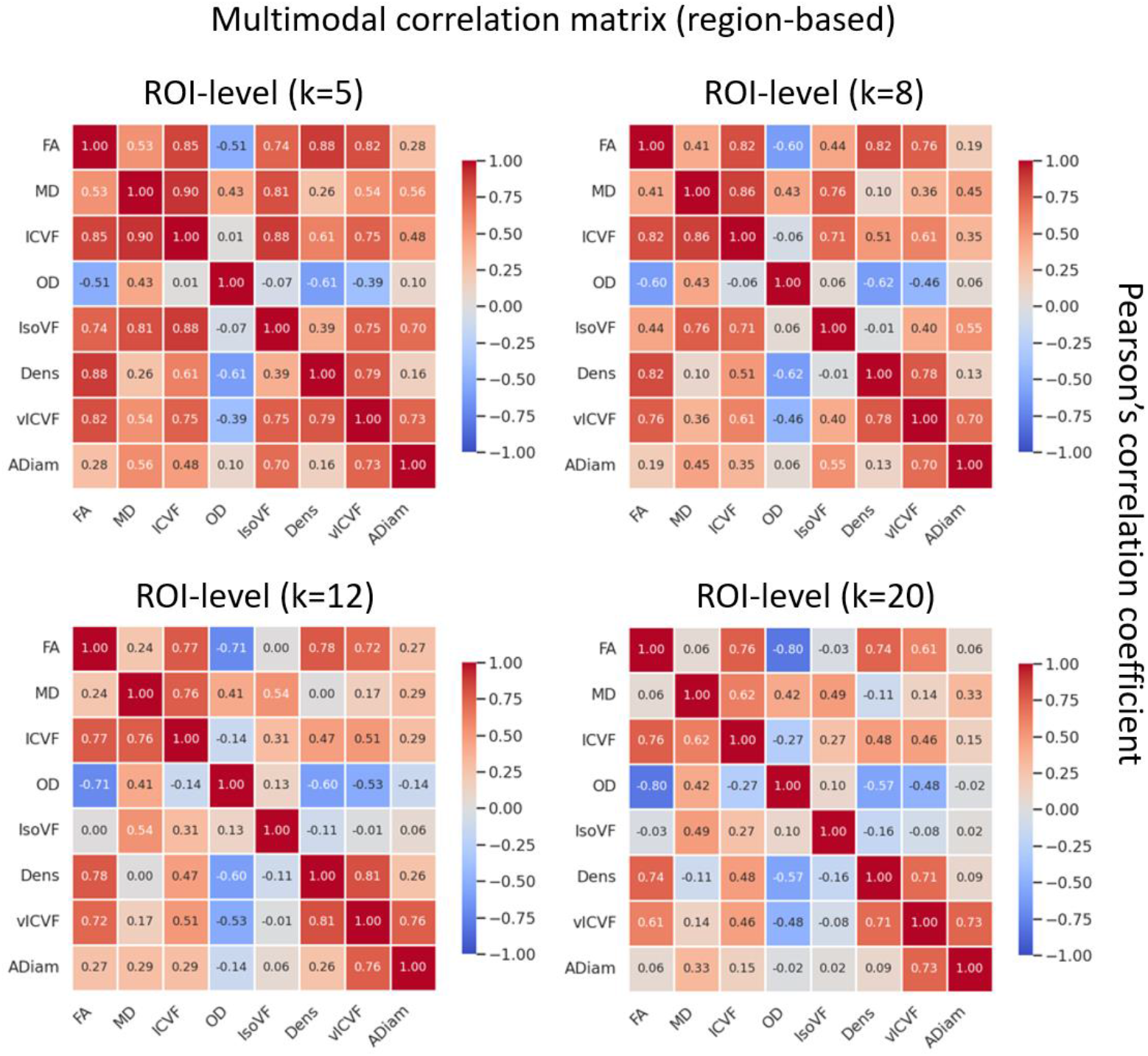
Scale-invariant multimodal correlation structure across region-based parcellations. Region-based Pearson correlation matrices are shown for diffusion-derived metrics computed after k-means parcellation of the brain into k = 5, 8, 12, and 20 regions of interest (ROIs). Despite changes in spatial granularity, consistent patterns of multimodal covariation are preserved across scales, including strong associations among axonal and intra-neuritic measures and antagonistic relationships involving orientation dispersion. These results demonstrate that the multimodal correlation structure is robust to spatial aggregation, supporting a scale-invariant organization of diffusion-derived metrics.

At coarse parcellations (k = 5), strong positive correlations emerge among FA, ICVF, IsoVF, AAx_Dens, and AAx_vICVF, whereas OD shows consistent negative correlations with anisotropy- and density-related metrics. As parcellation granularity increases, the magnitude of individual correlations varies, but the relative grouping of metrics remains stable. Notably, the tight association between AAx_vICVF and AAx_ADiam persists across all parcellations, as does the antagonistic relationship between OD and FA.

At finer scales (k = 12 and k = 20), although local heterogeneity increases and some correlations weaken, the same clusters of metrics continue to emerge. This scale invariance indicates that the observed covariation structure is not driven by voxel-level noise or region-specific averaging effects, but reflects global organizational properties of cerebral microstructure.

### Low-dimensional organization of multimodal diffusion space

Across voxelwise and region-based analyses, the diffusion metrics consistently organize into a limited number of coherent groups. One cluster is dominated by axonal and intra-neuritic measures (ICVF, AAx_Dens, AAx_vICVF, AAx_ADiam), another by diffusivity and dispersion-related metrics (MD, OD, IsoVF), with FA occupying an intermediate position reflecting sensitivity to both axonal coherence and dispersion.

This reproducible clustering suggests that the high-dimensional space of diffusion-derived metrics is effectively constrained to a reduced number of dominant modes of variation. These modes are not specific to any single diffusion model, but instead emerge from the structured relationships among metrics derived from distinct physical assumptions. These results therefore demonstrate that multimodal diffusion MRI encodes a low-dimensional organization of cerebral microstructure, consistent with the presence of latent biophysical diffusion properties underlying the observed signal.

## Discussion

In the present study, rather than treating diffusion metrics as independent descriptors, we examined whether structured covariation among diffusion MRI metrics derived from distinct biophysical models reflects underlying organizational principles of cerebral microstructure. By focusing on the joint statistical structure of multimodal diffusion measures rather than on isolated parameter maps, we demonstrate that diffusion-derived metrics consistently organize into stable, low-dimensional patterns across spatial scales. These findings support the interpretation that multimodal diffusion MRI encodes latent biophysical diffusion properties underlying cerebral microstructure in the squirrel monkey.

### Multimodal covariation as evidence of constrained biophysical degrees of freedom

Diffusion MRI metrics are often treated as independent descriptors of microstructure, each associated with a specific biophysical model or compartmental interpretation. However, as shown in Figures, metrics derived from tensor-based, multicompartment, and axonal models do not vary independently. Instead, they exhibit robust and reproducible patterns of covariation, suggesting that their variation is constrained by a reduced set of effective degrees of freedom.

This observation aligns with theoretical work in diffusion MRI demonstrating that many microstructural parameters are coupled through shared dependence on tissue geometry, compartmentalization, and restrictions imposed by cellular membranes (Novikov et al., 2018; Novikov et al., 2019). From this perspective, different diffusion models represent alternative parameterizations of a common underlying physical system, rather than independent probes of unrelated properties. The strong clustering observed among axonal and intra-neuritic metrics, as well as the antagonistic relationships between anisotropy and dispersion-related measures, are consistent with this view.

### Latent biophysical diffusion properties as emergent tissue states

The low-dimensional organization revealed by multimodal covariation suggests the existence of latent biophysical diffusion properties—effective tissue states that are not directly observable through any single diffusion metric, but emerge from the structured relationships among multiple observables. Similar concepts have been proposed in the context of “effective parameters” in diffusion physics, where complex microstructural environments are described by reduced representations that capture dominant constraints on molecular motion (Callaghan, 1991; Kiselev, 2017).

In neural tissue, such latent properties may reflect combined effects of axonal packing density, caliber distributions, orientational coherence, extracellular space, and membrane permeability. Importantly, these properties need not correspond to uniquely identifiable cellular features; rather, they represent stable configurations of the tissue that govern diffusion behavior at the mesoscopic scale accessible to MRI. The persistence of these covariation patterns across voxelwise and region-based analyses further supports their interpretation as global organizational features rather than local anatomical artifacts.

### Relation to neurodevelopmental organization and structural stability

From a developmental perspective, the emergence of stable latent diffusion properties is consistent with models in which cerebral microstructure results from dynamic neurodevelopmental processes operating under strong physical and geometric constraints. Theoretical and experimental studies have shown that axonal growth, cortical folding, and connectivity patterns are shaped by mechanical tension, material properties, and optimization principles that reduce degrees of freedom over developmental time (Van Essen, 1997; Budday et al., 2015).

Within this framework, the adult microstructural organization observed through diffusion MRI can be interpreted as a steady-state outcome of these developmental dynamics. The scale-invariant covariation patterns reported here suggest that the same underlying constraints operate across multiple spatial levels, from local microstructure to larger anatomical regions. This interpretation resonates with systems neuroscience views in which macroscopic brain organization emerges from the stabilization of multiscale interactions among cellular, geometric, and physical factors (Innocenti & Price, 2005).

### Implications for interpretation of diffusion MRI metrics

The present findings have important implications for how diffusion MRI data are interpreted. Rather than attributing biological meaning to individual diffusion metrics in isolation, our results support an integrative approach in which meaning arises from the relationships among metrics. This perspective helps reconcile longstanding debates regarding parameter specificity and identifiability in diffusion MRI (Jones et al., 2013; Novikov et al., 2018), emphasizing that diffusion metrics are best understood as coupled observables of an underlying biophysical system.

By framing multimodal diffusion MRI as a projection of a low-dimensional biophysical state space, this work provides a conceptual basis for reducing redundancy, improving interpretability, and guiding model selection in future studies. Importantly, this framework does not require assuming that any single model provides a “ground truth” representation of tissue microstructure.

### Limitations and future directions

Several limitations should be acknowledged. First, although the covariation patterns are robust, diffusion MRI remains an indirect measure of tissue microstructure, and the latent properties inferred here cannot be uniquely mapped to specific cellular mechanisms. Second, the analysis was performed on adult animals, precluding direct assessment of how these latent dimensions emerge during development. Longitudinal developmental datasets would be required to directly test the hypothesized link between neurodevelopmental dynamics and the stabilization of diffusion properties.

Future work may extend this framework by integrating multimodal diffusion covariation with histological validation, biomechanical modeling, or developmental trajectories. Additionally, dimensionality reduction techniques explicitly designed to extract latent variables could be used to further formalize the structure suggested by the correlation analyses presented here.

## Conclusions

In summary, this study demonstrates that multimodal diffusion MRI metrics exhibit structured, scale-invariant covariation reflecting a low-dimensional organization of cerebral microstructure in the squirrel monkey. These findings support the interpretation that diffusion MRI encodes latent biophysical diffusion properties arising from constrained tissue organization, rather than independent descriptors tied to individual models. By emphasizing the emergent structure of multimodal relationships, this work advances a systems-level, biophysically grounded framework for interpreting diffusion MRI in primate brains.

## Acknowledgments

The authors thank the National Council for Scientific and Technological Development (CNPq) and the staff of the Disconnectome Project.

